# Simulating Interclonal Interactions in Diffuse Large B-Cell Lymphoma

**DOI:** 10.1101/2023.09.28.559950

**Authors:** Siddarth Ganesh, Charles M. Roth, Biju Parekkadan

## Abstract

Diffuse large B-cell lymphoma (DLBCL) is one of the most common types of cancers, accounting for 37% of B-cell tumors globally. DLBCL is known to be a heterogeneous disease, resulting in variable clinical presentations and the development of drug resistance. One underexplored aspect of drug resistance is the evolving dynamics between parental and drug-resistant clones with the same microenvironment. In this work, the effects of interclonal interactions between two cell populations - one sensitive to treatment and another resistant to treatment - on tumor growth behaviors were explored through a mathematical model. *In vitro* cultures of mixed DLBCL populations demonstrated cooperative interactions and revealed the need for modifying the model to account for complex interactions. Multiple best-fit models derived from *in vitro* data indicated a difference in steady-state behaviors based on therapy administrations in simulations. The model and methods may serve as a tool in understanding the behaviors of heterogeneous tumors and in identifying the optimal therapeutic regimen to eliminate cancer cell populations using computer-guided simulations.

**Importance:** The cellular makeup of tumors can play a vital role in its growth and cancer development. In this work, two different types of cell populations of diffuse large B-cell lymphoma (DLBCL) were studied together to understand how they interact with each other in cultures. In mixed cultures, both types of cells cooperated with each other and increased their growth in complex manners. A mathematical model was created to simulate the growth behavior of mixed cultures. The model can potentially be used to predict future cell behavior and help in identifying more effective therapy regimens to maximize tumor cell reduction.

## Introduction

Diffuse large B-cell lymphoma (DLBCL) is one of the most common types of lymphoid malignancies, accounting for 37% of B-cell tumors globally with an estimated annual incidence of 15-20 per 100,000 in Europe and the USA [1]. DLBCL is known to be a heterogeneous disease, resulting in variable pathological and clinical presentations. Therefore, it has proven necessary to modify standard cancer therapies to combat the aggressive course of DLBCL. One of the most common treatments for patients with DLBCL is the combination of the anti-CD20 antibody rituximab (R) alongside the small molecule drugs cyclophosphamide (C), doxorubicin (hydroxydaunorubicin, H), vincristine (Onvocin, O), and prednisone (P). The R-CHOP regimen has considerably improved patient outcomes, resulting in disease-free rates between 50% to 70% [2]. However, this leaves 30% to 50% who will not be cured through this treatment, as 20% of the patients relapse after the entire treatment and 30% relapse in the middle of the regimen. About 80% of relapsed patients will die due to lymphoma, even after implementing therapies such as salvage chemotherapy or stem cell transplants [3]. It is therefore important to understand the mechanisms governing DLBCL development and relapse to create more effective treatment regimens and improve patient outcomes.

DLBCL drug resistance is thought to be caused by multiple factors, including the cell of origin, tumor microenvironment, variabilities between patients, and clonal evolution [3] [4]. A clone is a group of cancer cells that share a similar genetic profile. This clone can evolve over time through mutations and natural selection, leading to the development of a heterogeneous population of cells with diverse genetic profiles. The increased genetic diversity can lead to the acquisition of new traits such as drug resistance, invasiveness, and immune evasion. The mixed population can also lead to clonal competition, where the mutations result in a difference in growth behavior and prevalence. Clonal equilibrium, where subclone proportions are stably maintained in the tumor [5], is of interest to target and destroy all relevant cells within a tumor microenvironment. In clinical cases, DLBCL tumor samples contain significant heterogeneity, with clonal evolution observed in relapsed scenarios [6, 7]. The continuation of cooperative functions between the two types of cells, potentially maintained by the secretion of signaling molecules, nutrients, and other diffusible factors, could result in increased drug resistance or metastatic properties. Cooperation between clones has been observed in multiple cancer types other than DLBCL. In breast cancer, tumors composed of basal and luminal genotypes can be generated with increased secretion of the signaling molecule Wnt1, and they can evolve to rescue Wnt pathway activation in the molecule’s absence [8]. Non-genetic cooperative adaptation to therapy has been demonstrated in heterogeneous non-small-cell lung cancer tumors and has reduced sensitivity to radiation in prostate cancer [9, 10].

In the present study, a mathematical model was created to track cellular growth accounting for any reciprocal interactions between genetically distinct cell lines that can be sensitive or resistant to therapy by tracking the differences in steady-state responses with variations of initial conditions. *In vitro* studies utilizing DLBCL cell lines that were sensitive or resistant to the antibody Obinutuzumab validated the mutualistic interaction theory, though highlighted an imbalanced weighting of interaction between the drug sensitive and resistant cells. The mathematical model was then modified with a weighting function and further studied to see simulated events under different interaction weights. Simulations with weight-adapted growth rates were then developed using clinical growth rates parameters and also under drug pressure to explore steady state behaviors over long-term periods.

## Results

### Tumor Clones Exhibit Different Steady State Behaviors Based on Interactions

A mathematical model was developed that considers two cancer clones growing simultaneously, with a capacity constraint on both populations that reflects available space and/or nutrients. The logistic function was chosen for this purpose, as it has previously been used to describe a tumor in which proliferation is constrained by the available space [36]. As both clones grow in the same location, the increase in both populations results in a decrease in both rates. The total, maximum cell number reached is referred to as the carrying capacity. Initial monocultures of DLBCL cell lines demonstrated that drug-resistant cells grew at a faster rate than drug-sensitive cells. Thus, the sensitive cell growth rate, *k_G_*, was approximated to be 1 day^-1^, the resistant cell growth rate, *k_R_* was 0.5 day^-1^, and the carrying capacity was 600,000 cells for the simulations based on those cultures.

First, the growth of the two populations with a shared carrying capacity but otherwise no interactions was simulated (Equations 1-2). In this case, the larger growth constant of the sensitive population resulted in more proliferation for the first 6 days **(Fig 1A)**. Both populations ultimately reached steady states influenced by both the carrying capacity and individual growth rates - the total number of cells on day 10 approached the carrying capacity of 600,000, and the sensitive steady state was three times larger than the resistant one. A phase portrait depiction with sample trajectories and different initial proportions more clearly describes the distinct steady states resulting from distinct initial proportions **(Fig 1B)**. The mixed cultures deviated towards the sensitive cell number axis due to the population’s larger growth rate constant. The total cell number of all the cultures, including the pure and mixed variants, was equal to the carrying capacity.

**Figure 1:**
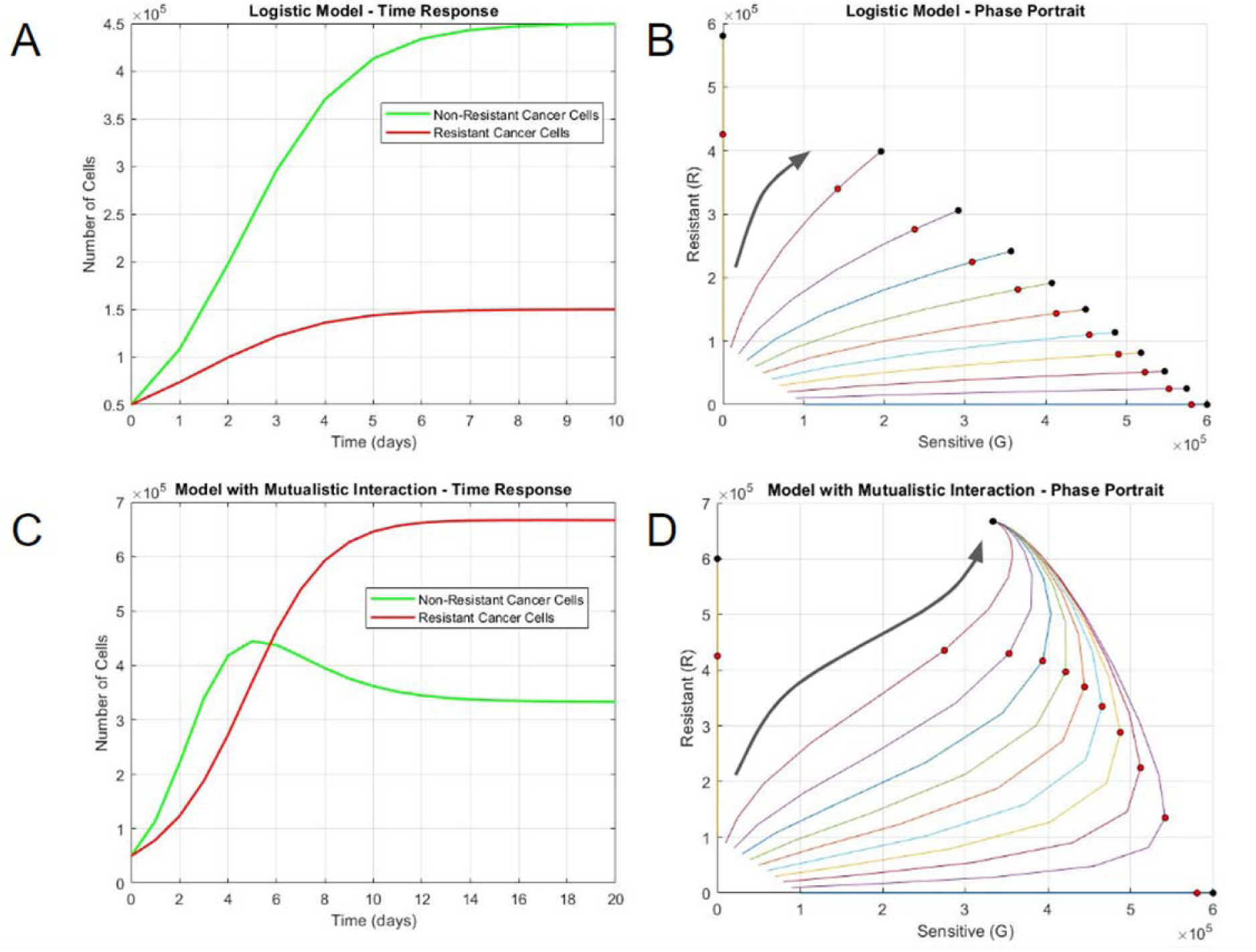
Long-term growth behaviors differentiate non-interacting and interacting cell populations. A) Time response and B) phase portrait of logistic model without interaction. C) Time response and D) phase portrait of logistic model with mutualistic interaction. In the phase portraits, the horizontal axis represents the number of sensitive DLBCL cells and the vertical axis represents the number of resistant DLBCL cells. Each curve in the phase portrait was created by altering the percentage of sensitive and resistant cells while keeping a constant initial population. The larger black arrows represent the general trajectories of population growth. The red circles represent the states on day 5, the black circle represent the steady state(s) at much longer time periods (day 100).

Next, the mathematical model was modified to incorporate potential interactions that may occur between the two DLBCL clones as evidenced in other cancers (Equations 3-4). Mutualistic interactions were simulated with positive interaction constants (*C_GR_* = 10^-6^ cell^-1^ day^-1^ and *C_RG_* = 10^-6^ cell^-1^ day^-1^). As the total population neared the carrying capacity, the mutualistic interactions resulted in a different steady state – resistant cells became the dominant population, as opposed to sensitive cells where mixed cultures always approached the steady-state solution, regardless of the initial concentration (**Fig 1C)**. This is even more apparent with the phase portrait, where mixed cultures always approached the steady-state solution, regardless of the initial concentration **(Fig 1D)**. Interestingly, the total cell number for the steady-state solution was more than the carrying capacity due to the interaction expressions in the system. While the system can yield steady-state solutions, increasing the interaction constants dramatically may result in solutions approaching infinity.

Competitive interactions were simulated with negative interaction constants (*C_GR_* = −10^-7^ cell^-1^ day^-1^ and *C_RG_* = −10^-7^ cell^-1^ day^-1^). Competitive interactions yielded two distinct steady-state solutions. Mixed cultures diverged towards the pure culture steady-state solutions depending on the proportion of sensitive cells and resistant cells **(Fig S1C, S1D)**. Parasitic interactions were simulated with one positive and one negative interaction constant (*C_GR_* = −10^-7^ cell^-1^ day^-1^ and *C_RG_* = 10^-7^ cell^-1^ day^-1^), to ensure the resistant population has an advantage. The initial time response (until day 3) was similar to that of the competitive simulation, but the parasitic nature of resistant cells ultimately led to the extinction of the sensitive cell population **(Fig S1A)**. Mixed cultures with multiple ratios approached the favored population carrying capacity, regardless of the initial concentration of sensitive cells **(Fig S1B)**.

### Rate Constants Determine Steady State Behaviors at Earlier Time Points

The steady-state behavior of mixed populations was most apparent at very large time points. Yet, there is a need to quantitatively describe the long-term behaviors of the different interactions to be able to compare them to *in vitro* results. The time responses of interacting and non-interacting systems cannot be analytically defined using the logistic function, as the cell numbers from both populations contribute to reaching the carrying capacity and the interaction constants provide deviations in initial behavior. Fitting the data from earlier time points (data until day 5) into a logistic equation was thus chosen as an approach to interpolate rate constants for direct analytical comparison between the simulations and *in vitro* results, since simulation results may not correlate with the total number fluorescent signals in potential in vitro studies. Fitting data points from each interacting and non-interacting simulation reveals a difference in growth rate constants **(Fig S2)**.

In the non-interacting model, as the resistant cell percentage increases, the rate of sensitive cells decreased in an almost linear manner, and as the resistant cell percentage decreased, the rate of resistant cells increased in a similar fashion **(Fig 2A)**. On the other hand, the sensitive cell rates increased for mixed mutualistic cultures with small initial percentages of resistant cells, but they decreased for higher percentages **(Fig 2B)**. Resistant cell rates in mixed mutualistic cultures demonstrated a slight decrease and a subsequent slight increase as the resistant percentage decreased.

**Figure 2:**
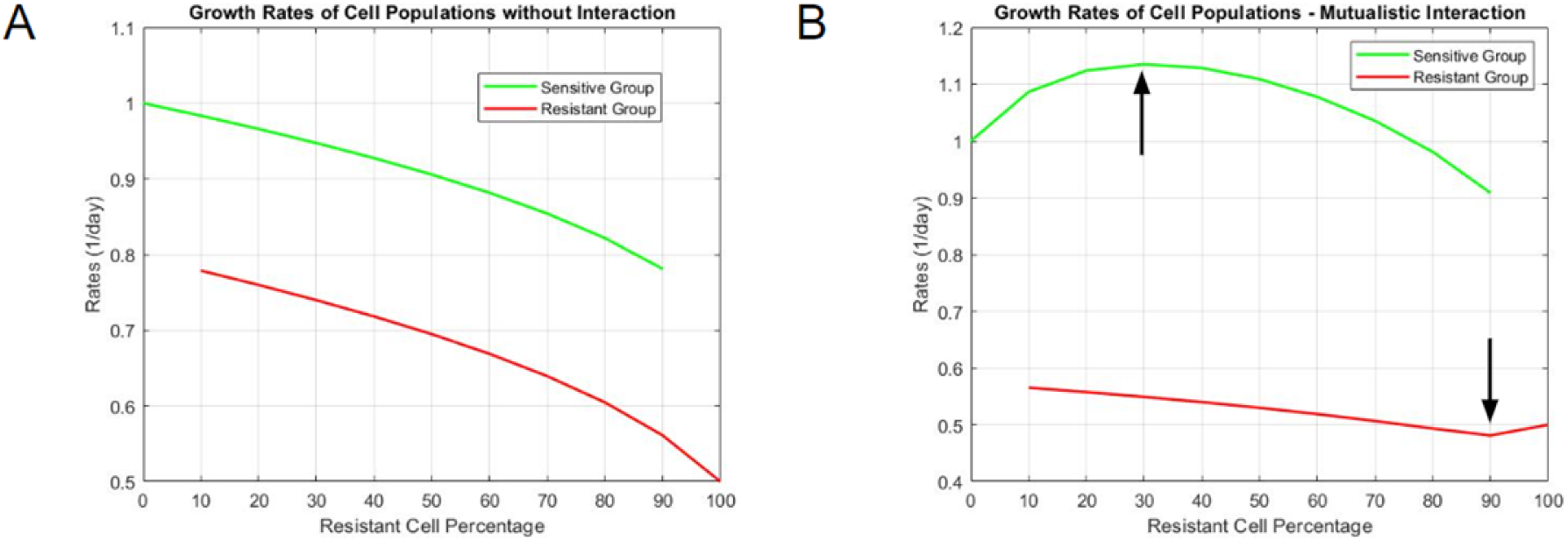
Interpolated growth rate constants for Systems A) Without interaction and B) With mutualistic interaction. The vertical axis represents the magnitude of rate constants, and the horizontal axis represents resistant cell percentage in each tested population proportion (a resistant cell percentage of 60 means the rate constants for each clone are derived from a starting culture of 40% sensitive DLBCL cells and 60% resistant DLBCL cells).

### In Vitro Cultures Exhibit Mutualistic Interactions Between DLBCL Clones

To study an interaction model with cell-specific precision experimentally, SUDHL-4 (drug-sensitive clones) and SUDHL-4OR (drug-resistant clones) cell lines were engineered to express green fluorescent protein (GFP) and red fluorescent protein (RFP), respectively. The cell lines were then grown as pure or mixed cultures. The cultures demonstrated logistic-like growth of both pure and mixed variants with the 3:1 and the 1:3 GFP:RFP ratio groups **(Fig S3)**. The primary periods of growth were from day 1 to day 4, and then the cultures approached a steady state on day 5. The G/R ratio was computed by dividing the GFP count by the RFP count. The ratios for the mixed cultures increased until day 2, indicating an initial increase of drug-sensitive cells **(Fig 3A)**. From day 3 to day 5, the ratio began to stabilize for the 1:3 GFP:RFP group but decreased for the other two groups, potentially indicating an interaction in which both populations cooperate, leading to one final steady state. Similarly, in simulations with mutualistic interactions, the G/R ratio started to converge after an initial distinction between each culture, regardless of their starting proportion of sensitive and resistant cells **(Fig 3B)**.

**Figure 3:**
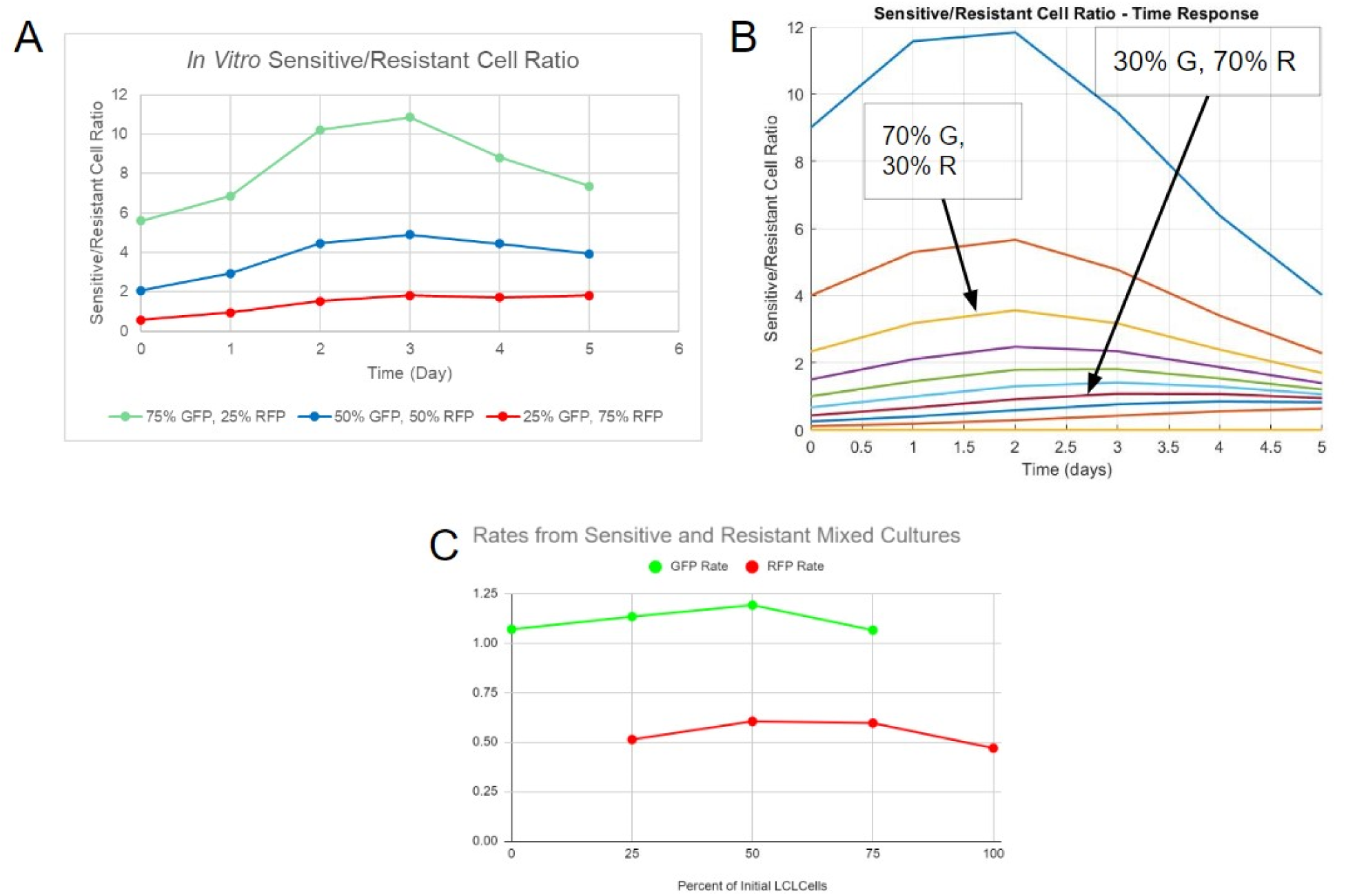
Data from *in vitro* study indicates mutualistic interactions between DLBCL clones. **A)** Ratios of sensitive/resistant cells of three tested proportions from *in vitro* fluorescence data. **B)** Ratios of sensitive/resistant cells of multiple proportions over time from simulation. **C)** Interpolated growth rate constants derived from fitting fluorescence data with the standard logistic function.

For the GFP count data, the rate constants slightly increased as the percent of initial resistant cells increases, with a maximum of 1.191 day^-1^ at 50% resistant cells **(Fig 3C)**. The rate constants of mixed populations were higher than the rate of pure sensitive cell culture (1.069 day^-1^). In regards to RFP count, the rate increased starting from the pure resistant culture (0.4695 day^-1^) as the percent of initial resistant cells decreased, with a maximum of 0.6045 day^-1^ at 50% resistant cells. Similar to its GFP counterpart, the RFP rates of mixed cultures were higher than the pure culture. The general behavior of GFP rates was most similar to that of the rates from mutualistic interaction simulations. However, the general behavior of the RFP rates did not match any rates from the simulations - it was instead similar to the GFP cell rates observed *in vitro*, with an increase and a subsequent decrease as the percent of initial resistant cells decreased. Although a mutualistic interaction was observed, a more complicated mode of interaction was likely needed to account for the disparity between *in vitro* results and simulations, and the similarity between the GFP and RFP cell rates *in vitro*. The model would also need to be modified to account for the steady-state cell number surpassing the carrying capacity in the initial simulations.

### Multiple Best Fit Models Derived Through Correlation

Since the first generation mutualistic model differentially captured the trends in rate constants with initial cell ratio for sensitive (GFP) vs. resistant (RFP) populations, a second generation model was constructed that would weight the interactions between the two cell types. This “weighting function” included a variable exponent (*m*) to the current sensitive cell number (*G*) to the conventional interaction expression and was divided by the total current cell number raised to the same exponent (*[G+R]^m^*) (Equations 5 and 6). The exponent will determine the maximum effect of interaction based on the starting proportion of cells. For example, when m is equal to 1, the interaction is maximized with 50% of the initial concentration of resistant cells, and this form is most similar to the conventional mutualistic interaction model form. However, as seen with the interpolated rates for that model, a maximum interaction at 50% does not directly result in a maximum interpolated rate at that same proportion. When m is equal to 2, the interaction is maximized with 33.3% of the initial concentration of resistant cells.

A weighting function was added to the interaction expressions in the differential equation system. Parameters were chosen based on the behaviors identified with *in vitro* cell growth. The sensitive cell growth rate constant, *k_G_* = 1.069 day^-1^, and the resistant cell growth rate constant, *k_R_* = 0.4695 day^-1^, were derived from interpolated rate constants from monocultures. The carrying capacity, *N* = 600,000, was approximated based on the total cell count that the cultures approached on day 5. Correlation tests were performed to derive interaction constants using different interaction exponents to represent in vitro data more accurately. A range of interpolated rate constants (similar to **Figure 2B**) were derived using a fixed exponent and a range of interaction constants *C_GR_* and *C_RG_*. Interpolated rate constants of sensitive or resistant cells were then correlated with their *in vitro* counterparts. The product of the independently derived correlation coefficients was maximized over the range of interaction constants.

**Figure S4** represents the correlation product when the two interaction constants are varied. The three-dimensional plot is indicative of the process used to obtain the constants - any possible combination of potential constants was checked for the correlation between the resultant simulations and *in vitro* data for a local maximum **(Fig S4A)**. The two-dimensional plot showed an example of how the correlation product compares to individual correlation constants of the sensitive and resistant groups **(Fig S4B)**. **Table S2** contains the different exponents used for the correlation test, along with their respective interaction constants and the correlation product. The correlation was lowest when the exponent equals one, which was closest to the conventional mutual interaction expression. Altering the exponent yielded higher correlation products. The value of the interaction constant C_GR_, representing the change in interaction for the growth of sensitive cells, increased based on the exponent. Fitting constants in this manner allows understanding how interaction exponents affect the final steady-states.

The steady-state solutions significantly varied with changes in exponents **(Fig 4A-D)**. Smaller interaction exponents resulted in the movement of steady states to the resistant cell axis, while larger exponents resulted in the dominance of sensitive cells. An exponent of 1 resulted in a similar proportion of final sensitive and resistant cells **(Fig 4E)**. Exponents between 0 and 1 yield larger final resistant cell counts. However, increasing the exponent from 1 resulted in increasing the final proportion of sensitive cells. All of the scenarios based on different interaction constant (*m*) values presented a possible model based on the analysis of interpolated rate constants and were carried forward for further simulations.

**Figure 4:**
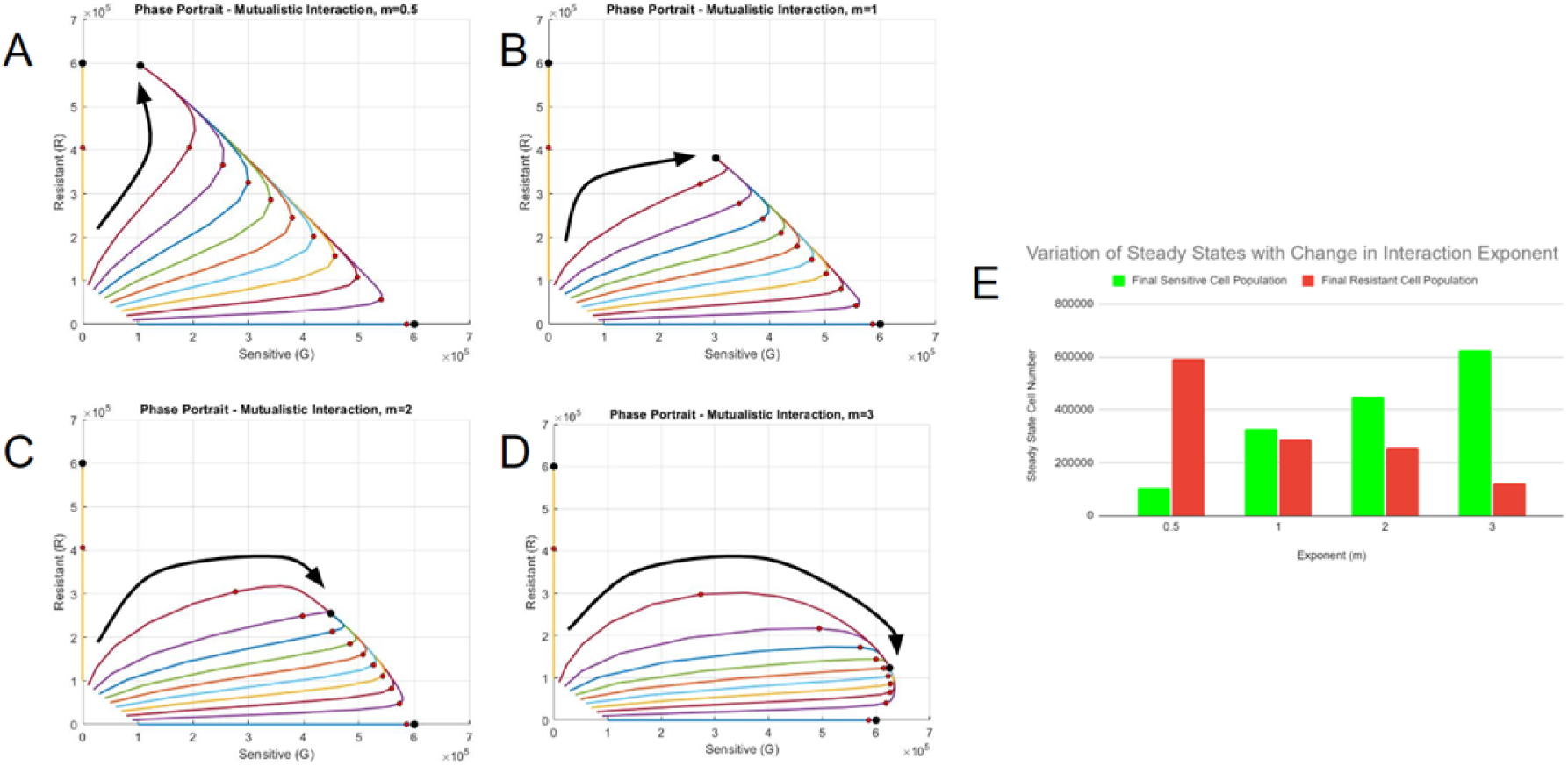
Correlation yields multiple best-fit models. Phase portraits of systems with best-fit constants with the interaction exponents A) m=0.5 B) m=1 C) m=2 D) m=3. The red circles represent the states on day 5, the black circle represent the steady state(s). The larger black arrows represent the general trajectories of each scenario. E) Final number of cells at steady state for each interaction exponent scenario.

### Best-Fit Models Exhibit Different Steady State Behaviors Based on Therapy Concentration

There have been strides to improve upon existing therapies to overcome therapy resistance in DLBCL. Rituximab, a chimeric IgG1 anti-CD20 antibody, has been effective in targeting CD20-positive B-cell malignancies through antibody-dependent cellular cytotoxicity (ADCC), complement-dependent cytotoxicity (CDC), and apoptosis usually referred to as “direct cell death” (DCD) [11]. However, cells can develop resistance to the antibody through multiple mechanisms, including downregulation or internalization of CD20, impairing ADCC through the production of C1q, and reducing CDC by the consumption of complement [12]. Obinutuzumab, a second-generation Fc-engineered anti-CD20 antibody, has been developed to overcome Rituximab resistance through the increase of ADCC and DCD capability [13, 11]. The antibody is thought to bind differently to B-cells than Rituximab, decreasing its CDC potency but invoking a greater DCD response that is caspase-independent [12]. However, the effects of the antibody may be affected by the interactions between cell populations within a heterogeneous tumor.

The impact of the Obinutuzumab therapy on the two interacting populations was tested via simulations to understand the relationship between cell growth and antibody-induced direct cell death. The direct cell death aspect of Obinutuzumab was added to the differential equation system through a time-varying function. A constant function of 1 was initially used to represent the constant presence of antibodies in the simulations. The interaction parameters with the exponent *m*=2 were initially used, with a starting population of 50,000 sensitive and 50,000 resistant cells, alongside an end time of 1000 days. The cell numbers at day 1000 and the phase portrait demonstrated two distinct steady-state solutions **(Fig 5A)**. At smaller concentrations of Obinutuzumab (0.3 or below), the culture approached a steady state which contained substantial amounts of both sensitive and resistant cells. The number of sensitive cells decreased as the therapy concentration increased to 0.3. This can also be seen in the phase portrait, in which distinct steady states (black dots) were achieved in the middle of the plot. However, with higher concentrations (greater than or equal to 0.31), there was an abrupt shift in the steady-state behavior **(Fig 5B)**. The number of sensitive cells at the end became zero, and the culture was dominated only by resistant cells. Moreover, increasing the concentration continued to affect the total cell number, as the number of resistant cells continued to decrease in a steady manner. This behavior was represented in the phase portrait with the solutions initially increasing to the right with sensitive cells with a subsequent reversal in behavior as they approached the resistant cell axis **(Fig 5A)**.

**Figure 5:**
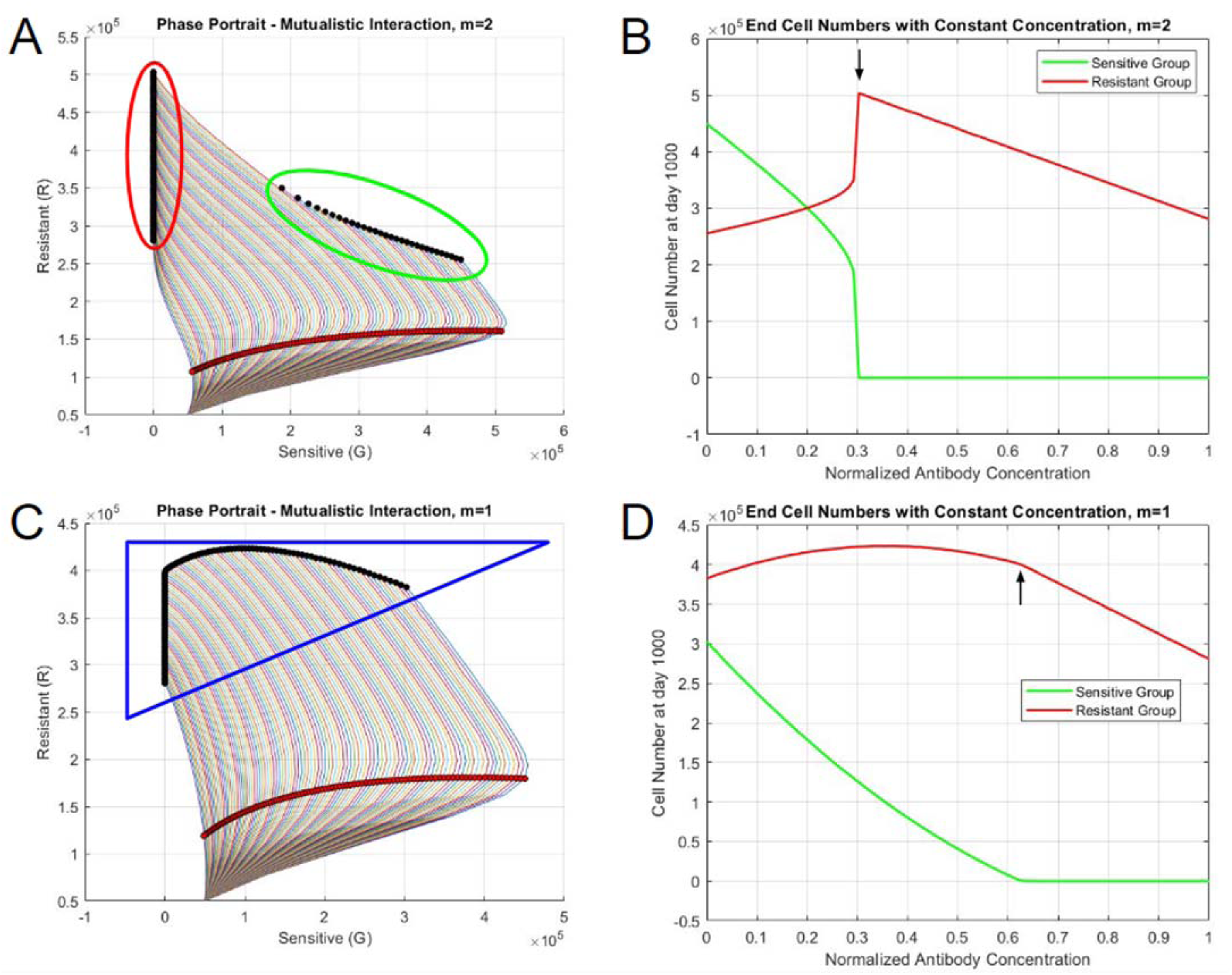
Steady state behaviors of systems with constant therapy. A) Phase portrait when m=2, B) End cell numbers when m=2, C) Phase portrait when m=1, D) End cell numbers when m=1. The red circles represent the states on day 5, the black circles represent the steady states. The outlines encircling sets of steady states represent distinct behaviors based on the concentration of therapy.

The critical point in steady-state was a function of the cooperativity parameter, m, as well as the normalized antibody concentration. When *m*=1, the abrupt shift did not occur. Rather, the number of sensitive cells steadily approached 0 as the antibody concentration increased until the concentration equals 0.63 **(Fig 5C-D)**. The resistant cell population only started to decline after the depletion of sensitive cells. When *m*=3, the change in steady-state behaviors occurred when the concentration is 0.41. The general behavior of the end cell numbers was similar to the ones from *m*=2, but there was a more sudden decrease in the sensitive cell number during the shift due to the large initial presence based on the interaction parameters and the exponent **(Fig S5A-B)**. When *m*=0.5, the general behavior was similar to when the exponent equaled to 1, but the resistant cell population continuously decreased regardless of concentration **(Fig S5C-D)**. The differences between the abrupt and non-abrupt shifts could also be identified in the phase portraits. The simulations with abrupt shifts yielded two separate clusters of final steady states, while the steady state solutions form the non-abrupt shift scenario were continuous in nature. The cluster formation seemed to be dependent on the exponent - two distinct groups form when the exponent is greater than 1.

In summary, based on variations of therapy concentrations and the exponent term in the interaction expression of the differential equation system, the scenarios tended to act in two different manners. Scenarios with exponents greater than one tend to abruptly change in behavior from a steady state containing substantial sensitive cells to one with only resistant cells with a small change in drug concentration, resulting in two distinct patterns. Scenarios with exponents lower than or equal to one tend to have non-abrupt decreases in sensitive cells with higher drug concentrations.

### Modeling Heterogeneous Tumors *In Vivo* with Clinical Parameters

In order to extend the previous simulations, which featured constants related to cell growth and interactions that were primarily derived from *in vitro* studies, both the constants and the manner of drug administration must be modified to simulate the clinical impact of interactions and therapy in vivo more accurately. More realistic cases of DLBCL were modeled by utilizing information from clinical tumor growth. It was noted that the doubling time of malignant lymphoma is approximately 29 days, which translates to a tumor growth rate of 0.0239 day^-1^ [14]. Furthermore, to assess the behavior of tumors with faster growth, a doubling time of 14 days was also considered as a separate case, which results in a growth rate of 0.0495 day^-1^. The size of observable tumors at the time of diagnosis was assumed to be 10^9^ cells, which would be the initial population of cancer cells in the simulation [15]. Different to the simulations above, a 9:1 ratio of sensitive and resistant cells would comprise the initial population (i.e., 9 x 10^8^ sensitive cells and 10^8^ resistant cells) in order to more realistically represent a human tumor population. The carrying capacity of a tumor within the body was approximated to be 2^40^ cells, which would equal to approximately 1 x 10^12^ cells [16]. In this case, it was assumed that the given growth rates correspond to a homogeneous sensitive cell population. Thus, the other constants (resistant cell growth rate constant and interaction constants) were scaled down to clinical constants. The list of constants is presented in Table S3.

A realistic dose application was used to depict a clinical course of antibody treatment for patients with DLBCL. The concentration of a therapeutic decays *in vivo* over time as drug molecules are cleared by the body. The kinetics of a finite amount of drug administered intravenously was governed by an exponential decay function, assuming that the pharmacokinetics are described by a single-compartment pharmacokinetic model. This function took into account the initial (maximum) concentration of the antibody, and the elimination constant (which modulates the speed at which the antibodies are removed from the system). In the case of a 1000 mg dose of Obinutuzumab, the maximum concentration of the antibody was 3.76 μmol/L and its half-life was 28.4 days [17]. The elimination constant, derived using the half-life, was 0.02441 day^-1^. In a clinical trial assessing the effectiveness of Obinutuzumab, patients were given IV administrations of the antibody in 21-day cycles [18]. During cycles 2-8, patients received 1000 mg of Obinutuzumab at the start of each cycle. During the first cycle, they instead were given 1000 mg injections on day 1, day 8, and day 15 (corresponding to 0, 7, and 14 in the simulation as the time starters at day 0). The plasma concentration of Obinutuzumab over the course of the clinical treatment with its clearance rate is shown in **Figure S6**, with a normalized concentration of 1 representing 1000 mg doses. Three initial administrations of the antibody were given to quickly increase plasma concentration, while subsequent administrations helped in maintaining a concentration range of approximately 1.5 to 2.25 times the initial dose. The concentration persisted until 400 days after the treatment was stopped on day 168. Obinutuzumab was administered alongside a regimen of CHOP that follows a similar 21-day cycle for 6 to 8 cycles. Most of the drugs present in CHOP (cyclophosphamide, doxorubicin, and vincristine) were infused via IV on day 1 of the cycle, while prednisone was given orally on days 1 to 5. Simulations were run for 1500 days, equaling approximately 4 years, with the start of the therapy set to day 0.

Both interaction tumor simulations end at a similar concentration to that of the non-interacting control, which instead demonstrated a lack of resistant cells at lower concentrations and a maximum of them at around 0.075 **(Fig 6A)**. In the simulation performed with slow-growing tumor constants, the behavior of total end cell numbers was consistent with both interaction exponents - tumor cells persist until the concentration reaches 0.12 **(Figs 6B-C)**. However, the difference was more apparent when the clone populations were considered. When *m*=1, the resistant cell population reached a peak around the concentration of 0.015 but continuously decayed until 0.12 **(Fig 6C)**. When *m*=2, it had two distinct maxima (when the concentration is 0.01 and 0.07) followed by a subsequent decay **(Fig 6B)**. A large distinction can be observed between fast-growing tumors. In terms of similarity, the control and both interaction simulations resulted in the cessation of tumor cells by the 0.3 concentration. However, the interaction simulations yielded persistent presence of the sensitive cells until certain antibody concentration thresholds, with the proportion of each clone dependent on the exponent. When *m*=2, the sensitive cells become depleted around the same concentration as the control simulation. Moreover, it demonstrated an abrupt shift in population proportion around that point when compared to the other scenarios. When *m*=1, however, both clones persisted for a longer concentration range until the sensitive variant gradually decreased, resulting in a peak of resistant cell population around 0.2.

**Figure 6:**
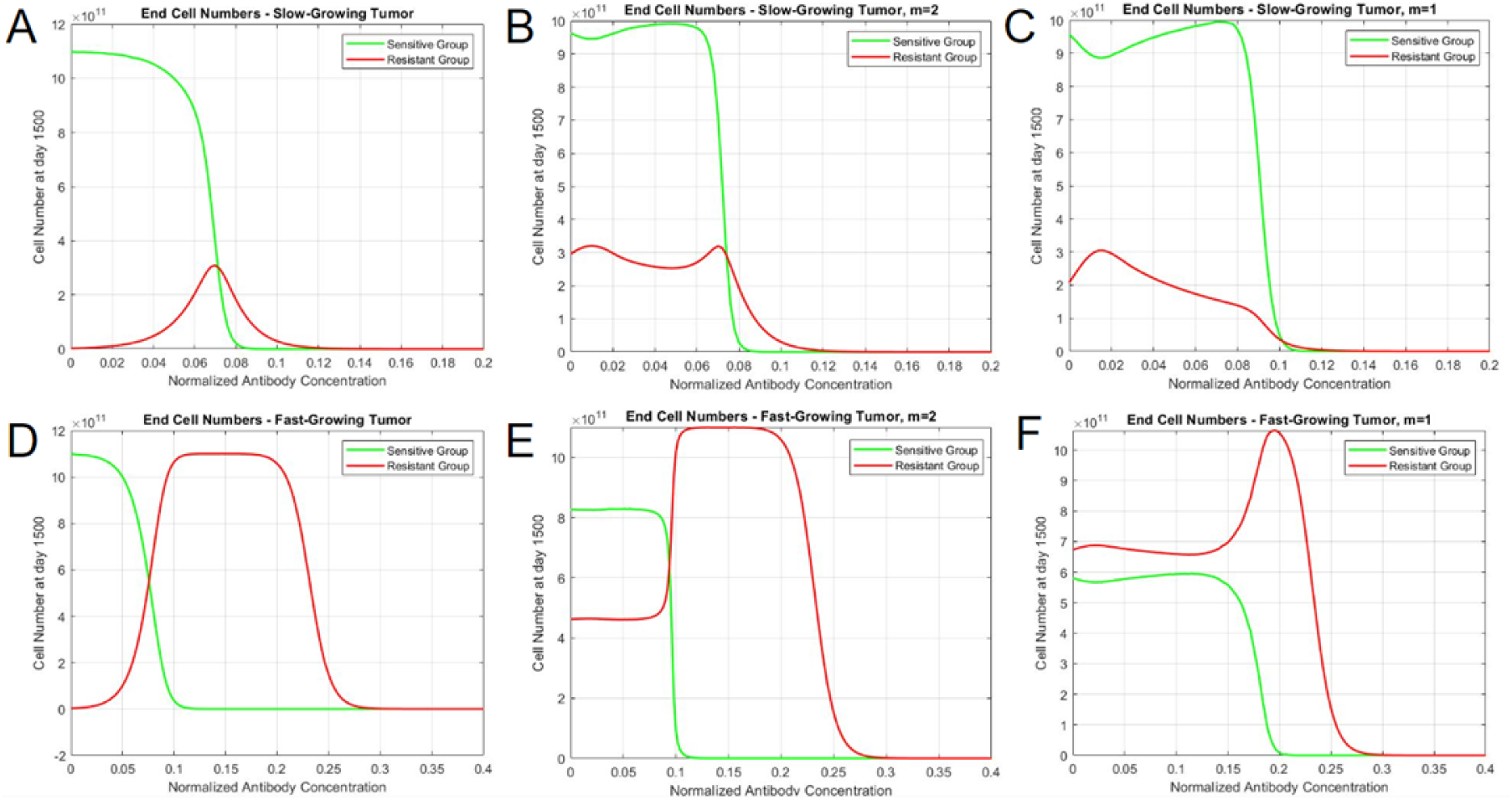
Tumor composition varies based on interaction type, tumor growth, and therapy concentration. End cell numbers for A) Slow-growing tumor, without interaction B) Slow-growing tumor, m=2 C) Slow-growing tumor, m=1 D) Fast-growing tumor, without interaction E) Fast-growing tumor, m=2 F) Fast-growing tumor, m=1. The vertical axis represents the cell number at day 1500, and the horizontal axis represents the magnitude of antibody concentration with a constant treatment cycle.

## Discussion

In this work, a model depicting interactions between different DLBCL cells was developed to understand the long-term behaviors of heterogeneous tumors. As a highly heterogeneous condition, DLBCL could lead to variable clinical presentations and treatment resistance. This work explored the effect of mixed populations and therapy regimens on the steady states. Interpolated rate constants were used to distinguish different types of possible interactions occurring in smaller timescales. Cells sensitive and resistant to therapy were engineered to produce fluorescent markers and were cultured as mixed populations in order to identify potential interactions. The G/R ratio additionally demonstrated an initial increase followed by a subsequent stabilization or decrease for groups with different initial cell proportions, suggesting a cooperative interaction. The interpolated rates of sensitive and resistant cells from *in vitro* growth deviate from the non-interacting and interacting simulations - the rate constants of both groups approached a maximum at 50% initial resistant cells. The interactions were also likely dependent on the initial proportion of sensitive and resistant cells due to the similarity in behavior between the sensitive and the resistant cell rates. As the *in vitro* rate constants differed from the previously derived constants, a modified interaction expression was added to the model.

Rate constant analysis at earlier time points was able to differentiate between interacting and non-interacting models. Tracking the mathematical differences in growth behaviors seen in mixed cultures can work as an alternative method to find interactions in cancer clones by comparing *in vitro* growth rate constants to interacting or non-interacting population simulations. Derivation of the interaction constants *CGR* and *CRG* was performed by correlating the *in vitro* interpolated rate constants with simulation derived by using a range of potential constants. Since the two constants are independently tested for, a product between both correlations was instead maximized in order to derive best-fit constants for each exponent. A similar method of multiplying coefficients is used in path analysis, a variation of multiple-regression analysis used to examine causal relationships between variables [45]. The correlation products increased with higher values of the cooperativity, *m*, supporting the introduction of this parameter into the model. Nonetheless, as the variation in correlation products was moderate, it was decided to continue with all four values of *m* for further analysis of scenarios involving therapeutic treatment.

Upon treatment with a drug that induces direct cell death, two different behaviors were identified in terms of the population steady-states, which depend on the interaction exponents. Some interaction exponents result in two distinct steady states, and others result in a singular steady state, as the therapy concentration increases. When the exponent was less than or equal to 1, the steady-state cell numbers for sensitive and resistant cells gradually decreased with higher therapy concentrations, resulting in a spectrum of steady states as observed in the phase portraits. However, exponents greater than 1 yield steady states with substantial proportions of sensitive and resistant cells with lower therapy concentrations, but abruptly shift to pure resistant cultures after surpassing a threshold drug concentration. Moreover, testing the model with *in vivo* parameters revealed variations in resistant cell population behaviors in slow-growing and fast-growing tumor cases. In terms of slow-growing tumors, both interaction types had similar tumor profiles throughout the concentration range, in which there was a mild persistence of resistant cells. In fast-growing tumors, the simulations demonstrated distinct behaviors based on interaction types. When *m*=2, both sensitive and resistant cells persist in an equilibrium until an abrupt shift occurs at the 0.99 concentration, which results in the depletion of green cells and an increase in red cells. In contrast, the simulation with no interaction only has sensitive cells at earlier concentrations, and the resistant cells become the dominant clone in a more gradual manner. When *m*=1, however, the resistant cells are dominant throughout the concentration range and approaches a maximum after the eventual depletion of sensitive cells.

Interactions between tumor cells and the surrounding non-tumor cells play an important role in promoting cancer cell proliferation or metastasis. In *in vitro studies*, mixed cultures tended to form free-floating clump-like structures which contained both sensitive (green) and resistant (red) cells. The process of clumping could be a potential mechanism through which interaction between the two types of cells occurs. In this work, we incorporated a non-capacity-constrained interaction term (Equations 3 and 4) to account for the possibility that the capacity of both space and nutrients is likely to be altered by interacting cell types. Nonetheless, a spatial component considering the distance between two cancer clones (such as a partial differential equation system) could more accurately depict their interactions within the spheroid-like structures that were observed here. Partial differential equations can be used in that case to understand the spatial variations that occur during tumor proliferation [19]. Multiple mechanisms could be involved in conferring advantages for tumor cells in heterogeneous clusters. Tumors could produce growth factors which can bind to their own surface or the surface of genetically similar clones, leading to enhanced proliferation due to autocrine signaling. Overexpression of epidermal growth factor receptor (EGFR) is known to enhance cell survival in epithelial cancers and gliomas [20] [21]. Moreover, cancer stem cells (CSC), a variant with stem cell-like properties, can interact with their non-CSC counterparts to enhance survival. Differentiated colorectal cancer cells have been known to protect the stem-cell variants from chemotherapeutic toxicity and glioma cells behaving like CSCs can produce IL-6 to promote growth of non-CSCs [22, 23].

To incorporate these types of interactions in simulations, a model would involve the inclusion of the genetically similar cell population with information regarding its growth and self-renewal capacity. Compartment models describing movements of cells between different types and competition models describing interactions between different types have been used to model the development of therapy resistance and heterogeneous tumors respectively [10] [24]. Moreover, it could also involve tracking the molecular kinetics of signaling molecules and receptor-binding processes. Paracrine signaling from other non-cancerous cells in the tumor microenvironment have also been known to enhance tumor growth. Stromal fibroblasts can promote tumor progression through remodeling the extracellular matrix and producing cytokines or transforming growth factor-β (TGF-β) [25, 26]. Cancer cells can also take advantage of immunomodulatory activities in the microenvironment to promote growth and metastasis. Cells expressing PD-L1 checkpoint can interact with PD-1 on T cells, leading to exhaustion and decreased antitumor responses [27]. Tumor-associated macrophages (TAMs) can lean towards the M2 phenotype and aid tumor cells through producing immunosuppressive cytokines and other factors to promote angiogenesis or chemotherapy resistance [25, 28, 29]. Macrophages can additionally influence phenotypic changes in cancer stem CSCs, leading to increased metastatic potential [46]. The inclusion of TAMs or effector cells would require the tracking of the noncancerous cell populations, which may require more incorporation of *in vivo* data as they are not self-renewing in the same capacity as cancer cells. Systems of ordinary differential equations consisting of tumor cells, antibody concentrations, and effector cells that attack cancer reflect the in vivo dynamics of therapies [14, 15].

The interaction expression in the mathematical model was modified with the additions of an exponent and a denominator in order to incorporate the in vitro observations. The denominator with the total population is included to ensure that the interaction constants will not fluctuate between large orders of magnitude when changing the exponent for testing the model. The denominator (*[G+R]^m^*) was added in order to reduce the deviation of constants in orders of magnitude observed while testing the modified model and account for the drastic increase of the populations over the carrying capacity seen in the initial simulations. It represents the total number of cells at a given time, and its inclusion ensures that the increase in cell growth is determined by the proportion of interacting cells. The “weighting” aspect of the function is an exponent placed with the sensitive cell population term, and its value ultimately determines the final steady states of the simulated cultures. The exponent is placed with the sensitive cell number rather than the resistant cell number since the alternative did not yield interpolated rate behavior similar to in vitro results. Biologically, the interaction exponent may represent when the cancer’s growth is maximized with a given tumor composition. For example, with an exponent of 2, a ratio of 2-to-1 sensitive-to-resistant cancer cells is required to maximize the interaction-based growth rate. When *in vitro* data was fitted using modified interaction expressions, exponents of 0.5, 1, 2, and 3 were used, which demonstrate requiring a near-equal number of sensitive and resistant SUDHL-10 cells to maximize interactions. However, different exponents and thus different interaction ratios may apply to other cancer types. A clinical ratio of cancer to stroma cells was used to predict the invasiveness of lung adenocarcinoma and in head and neck cancers [32, 33]. Effector-to-target cell ratios are also important at determining the effectiveness of the immune system against tumors, with a ratio of 100:1 maximizing effectiveness for neutrophil ADCC [34, 35]. The ratio of cells observed in proliferation or reduction are likely dependent on a number of factors, inducing the biochemical mechanisms involved in terms of molecular signaling and *in vivo* spatial constraints. A deeper exploration of the mechanistic aspects of DLBCL cell interactions would be required to understand the reason for a similar population of clones maximizing proliferation as opposed to larger ratios with tumor-non-tumor interactions.

The model was limited in scope in order to study potential interactions between two cancer populations. Multiple assumptions were used to describe initial cell growth and drug resistance. A logistic function was chosen to describe the general behavior of DLBCL cells, as it can be relevant when considering a spatially extended system in which proliferation is constrained by available space [36]. Moreover, logistic growth has been observed in clonal equilibrium associated with chronic lymphocytic leukemia, another hematological malignancy [37]. For initial computational simulations, it was assumed that the resistant cells would have a smaller rate constant than the sensitive variant. Drug-resistant cancer cells have been observed to grow at a slower rate, suggesting that reduced proliferating activity can contribute to resistance [38, 39]. While Obinutuzumab has multiple modes of action to induce apoptosis, ADCC requires the presence of effector immune cells to kill CD20-positive cells. Thus, for simplifying the current model, only the effects of direct cell death were analyzed instead of adding another cell population (effector cells). For the therapeutic effects of the antibody, it is assumed that the effect of the therapy is only dependent on the concentration of the molecules in the *f(t)* expression and not dependent on the effects of ligand-binding conformation changes that are characterized by the Hill equation. Assumptions regarding the potency were also made to simulate therapy effects. In one study, high concentrations of Obinutuzumab alone resulted in a proliferation percentage of 75%, starting from a percent of 100% without the antibody for an Obinutuzumab-resistant SUDHL-4 clone [13]. Assuming that the proliferation percentage for Obinutuzumab-sensitive cells would be 0% with a high antibody concentration, it can be interpreted that the therapy is one-fourth as effective against the resistant counterparts. Implementing other forms of therapy requires different mathematical expressions to depict their effects. The linear quadratic model incorporating tissue sensitivity constants is used to describe the effects of radiation therapy based on similar behaviors observed in kill curves [40]. Chemotherapy modeling, on the other hand, uses a similar expression to the one used in this work to describe antibody effects, in which the dose and population number decreased growth rate [41]. As both of the aforementioned treatment modalities are less specific, they would likely require less information on *in vivo* and microenvironment conditions to accurately predict responses. Since antibody therapy is more targeted, a more complex model would incorporate the effects of direct cell death with receptor binding kinetics and the effector cell populations (such as T cells or NK cells) with their respective affinities to the molecule for the effects of ADCC.

Further studies could provide more challenging tests of the assumptions in the mathematical model. An *in vitro* study can be performed with exploration of potential molecular interactions such as growth factors or cytokines to assess the assumptions made in the model. An experiment with variations in both culture heterogeneity and Obinutuzumab concentration can explore the protective effects of mixed tumors under therapeutic stress. Moreover, it can reveal potential abrupt changes in cell populations within specific concentration ranges, establishing which best-fit model would most accurately describe the interaction process. The use of genetic or epigenetic analyses and growth factors in cultures can aid in determining which biochemical pathways the cancer clones use for interactions. For example, methylation of TGF-β associated genes was linked to therapy relapse in DLBCL cells [42]. Thus, the use of TGF-β in pure and mixed cultures could reveal alteration of growth patterns indicative of a change in interactions. Moreover, parameters could be varied to accommodate other aspects of cancer growth and therapy, including factors in tumor microenvironment and the presence of effector cells.

The results of human-based studies in literature do not completely suggest the validity of a model with one interaction exponent over another. A primary method of studying the variations between regular and relapsed/refractory DLBCL is through mutational analyses of tumor biopsies. These studies have demonstrated that specific gene alterations necessary for immune surveillance and suppression are observed in higher frequency in samples after relapse, and some of them are correlated with reduced overall survival [13, 43, 44]. The patients would have likely been treated with therapies based on a constant dose or based on body weight, and analyses after relapse do not capture the dynamics involved between therapy-sensitive and therapy-resistant clones. Thus an *in vivo* study analyzing tumors at multiple time points, from a diagnostic size to the time of relapse (potentially by tracking general tumor growth behaviors via growth rate constant derivation) would give better understanding on which best-fit model would most efficiently capture real-life observations - an abrupt shift in clonal populations because of increasing therapy concentration would favor larger interaction constants.

The modified interaction model could be useful as a clinical predictor of heterogeneous tumor growth and aid in optimizing therapy regimens. The results indicate a need to evaluate interclonal interactions and use different approaches to reduce specific cell populations. While conventional therapy may be more necessary in clearing drug-sensitive cancer cells in the cases where the interaction exponent equals one, altering it to a different medication to coincide with the abrupt shift of cell populations noted with larger exponents could be more efficient in reducing tumor burden. Both the patients’ tumor growth datasets and mutation profiles can be used to assess the interaction exponents involved and predict future proliferation behavior. Therapy concentration and duration can subsequently be altered to be most effective during different phases of tumor growth. Combining predictive mathematical models with specific regimens enables physicians to offer personalized medicine to treat patients more effectively.

## Methods

### Simulation Parameters

All differential equation systems were simulated using MATLAB R2020a. The “ode45” and “ode23” functions were used to compute the changes in sensitive cell and resistant cell populations over time. The system consisted of a sensitive cell population (*G*) and a resistant cell population (*R*). The inputs for the system included the initial number of cells for each clone, the time (in days) for how long the simulation takes place, and a function representing the therapy concentration over time.

### Initial Simulations of Differential Equation System

A differential equation system based on the logistic growth model was used to model potential interactions that may occur between the two DLBCL clones. The Lotka-Volterra model, a system originally used to describe predator-prey population behavior, has been used to describe the interaction between prostate cancer cells [8=10]. The equations below include expressions needed for interactions, alongside the aforementioned logistic components. “*G*” or “sensitive cells” in the equation system represents the population consisting of DLBCL cells sensitive to therapy, and “*R*” or “resistant cells” represents the drug-resistant counterpart. The rate constants for each population and the carrying capacity are represented as “*k*” and as “*N*” respectively.

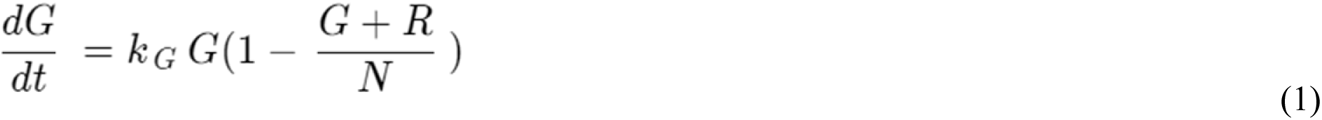

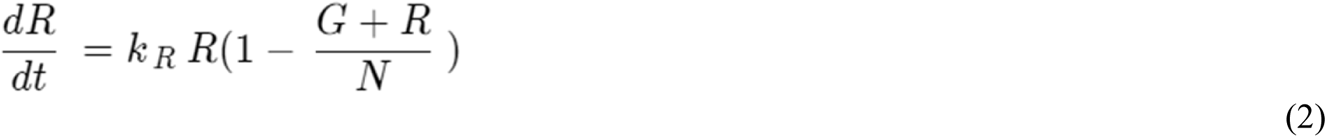

The logistic system was modified to incorporate theoretical interactions that may occur between the two DLBCL clones. The equations below include expressions needed for interactions, alongside the aforementioned logistic components.

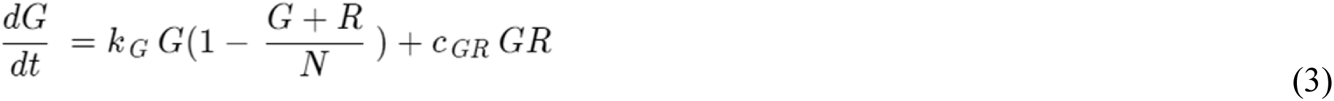

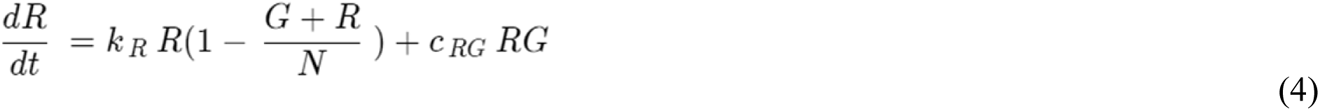

In this case, C_GR_ represents the interaction constant that alters the sensitive population growth rate, while C_RG_ represents the constant modifying the resistant cell growth, and the product between both clones mirrors the extent of interactions that may occur within cultures.

### Rate Constant Derivation

The rate constants were derived individually from sensitive and resistant cells in mixed cultures for each proportion (e.g. 100% sensitive cells, 100% resistant cells, 50% sensitive and 50% resistant cells), mimicking how the separate fluorescence data can be captured *in vitro*. The data points until day 5 from interacting and non-interacting simulations were inputted into MATLAB’s curve fitter application (cftool) individually and a regular logistic function was used to fit the data. The rate constant estimate from the algorithm was identified from each individual population (drug-sensitive or drug-resistant) and repeated over multiple proportions of each clone.

### DLBCL Cell Lines, Transfection, and Transduction

SUDHL-4 (sensitive cells), a DLBCL suspension cell line, and SUDHL-4OR (resistant cells), a DLBCL suspension cell line resistant to Obinutuzumab, were used for the study. The cells were obtained through the Rutgers Cancer Institute of New Jersey (CINJ). The cells were cultured in RPMI 1640 media (ATCC 30-2001) containing 4500 mg/L glucose supplemented with 10% fetal bovine serum (FBS) and 1% antibiotic-antimycotic solution (Fisher Scientific, Boston, MA) at 37 °C in a humidified atmosphere with 5% CO2. Cell viability and proliferation was quantified using automated NucleoCounter NC-202^TM^ with an aliquot of 200 μL.

Cells were engineered to produce distinct fluorescent proteins in order to track them for proliferation measurements. Genetic constructs (plasmids) that constitutively produce green fluorescent protein (GFP) and red fluorescent protein (RFP) were obtained through VectorBuilder. The plasmids are as follows: pLV-EF1α-GFP and pLV-EF1α-RFP. The plasmids were packaged in lentiviral particles through transfection. HEK293T cells were cultured in Opti-MEM media with PEI transfection reagent, a packaging plasmid (psPAX2), a viral envelope plasmid (pMD2.G), and each genetic construct in order to produce viral particles containing the constructs. A 3:2:1 ratio of the plasmid of interest, psPAX2, and pMD2.G was used. The media was replaced with RPMI 1640 24 hours after the initiation of the process. The remaining media was collected 48 hours after the initiation of the process and filtered using 20 mL syringes and 0.45 μm syringe filters to obtain viral particles. Concentrations of viral particles were verified via qPCR using a lentiviral titration kit from Applied Biological Materials on QuantStudio 3 from Thermo Fisher.

In the process of spinoculation, SUDHL-4 cells were centrifuged in RPMI 1640 media containing viral particles with GFP construct, and SUDHL-4OR cells were centrifuged in RPMI 1640 media containing viral particles with RFP construct, in order to transduce the cells for fluorescent protein expression. The cells were centrifuged at 2400 RPM for 2 hours and were subsequently kept inside the centrifuge for 2 hours. After culturing the transduced cells, some sensitive cells were able to express GFP and some resistant cells were able to express RFP. The two cell populations were sorted via flow cytometry to obtain pure fluorescent cells. The process resulted in SUDHL-4 cells fluorescing green due to GFP expression and SUDHL-4 OR cells fluorescing red due to RFP expression **(Fig S5)**.

### Mixed DLBCL Cultures

Pure and mixed initial cell populations with a total initial cell number of 100,000 in 1 mL RPMI 1640 media for each group were cultured in 24-well plates to simulate a spatial constraint. Different ratios of mixed populations were cultured in order to determine differences in rates between the cultures. Ratios of 1:0, 3:1, 1:1, 1:3, and 0:1 G:R cells were tested. The day on which the cells were initially seeded was defined as the 0th day for the time in experimentation to match with computer models. 0.5 mL media was added to each well on days 2 and 3 in order to maintain nutrition. The fluorescence and total cell count were determined each day using the Celigo imaging cytometer **(Fig S7)**. The Celigo imaging cytometer can collect information regarding cellular fluorescence by scanning well plates without the need to remove samples from each group for flow cytometry analysis.

While scanning plates, it was noted that both cell populations tended to clump together during prolonged periods of time. The clumps of cells contained both sensitive and resistant cells, determined by the presence of green and red fluorescence respectively **(Fig S7)**. In order to properly image the suspended cellular clumps, the cells were separated by mixing the cultures with a pipette and subsequently centrifuged so that they lay flat on the bottom of the well plate. The *in vitro* data was then analyzed using MATLAB’s curve fitter application (cftool) and a conventional logistic function to fit the data and derive constants for the rate (*k*), initial population (*b*), and the carrying capacity (*N*) **(table S1)**.

### Simulation - Differential Equation System with Modified Interaction Expression

The system of equations including the modified interaction expression (equations 8 and 9) is shown below. Parameters were chosen based on the behaviors identified with *in vitro* cell growth. The sensitive cell growth rate constant and the resistant cell growth rate constant were derived from interpolating constants. The carrying capacity was approximated based on the total cell count the cultures approached on day 5.

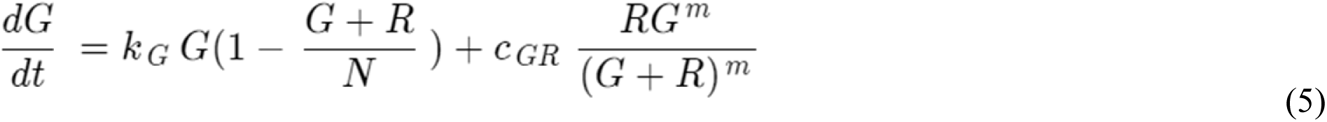

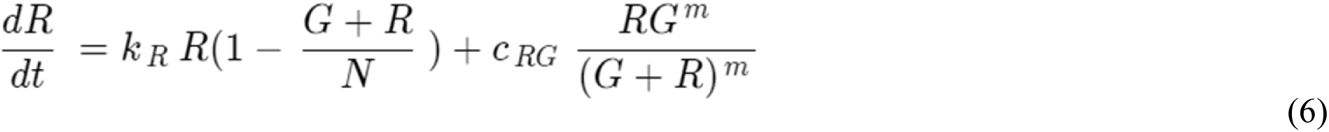

### Derivation of Interaction Constants Through Correlation

MATLAB’s “corrcoef” function enabled correlation with two separate sets of data. The function outputs a number close to 1 if the data sets (*in vitro* and simulation) are identical, 0 if they are not correlated, and −1 if they are inversely correlated. The correlation coefficients were derived through a comparison between the *in vitro* interaction constants and simulation interaction constants for both the sensitive cells and resistant cells. The interpolated rate constants of sensitive cells were correlated with its *in vitro* counterpart and the constants of resistant cells were separately correlated with its *in vitro* counterpart. In order to maximize the correlation of both populations at the same time, the product of their respective correlation coefficients was assessed. If the correlation product is close to 1, it would mean that both populations would be represented accurately by one set of interaction parameters.

### Simulation - Differential Equation System with Cytotoxic Antibody Drug Treatment

Equations were modified to incorporate the effects of direct cell death caused by Obinutuzumab. The constants *a_G_* and *a_R_*represent the antibody’s potency against the populations of sensitive and resistant cells respectively. The function *f(t)* represents the administration of the therapy with time as the independent variable. The function *f(t)* represents the function of therapy concentration over time. For the simulation, both potency constants were normalized so that *a_G_* would equal 1 and *a_R_* would equal 1/4.

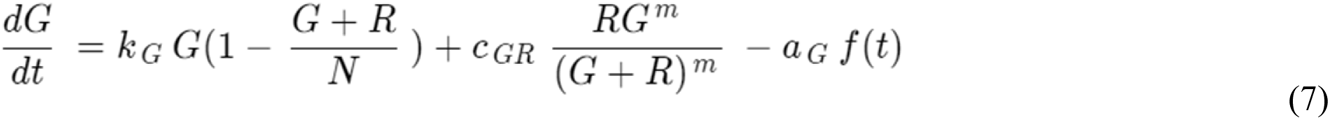

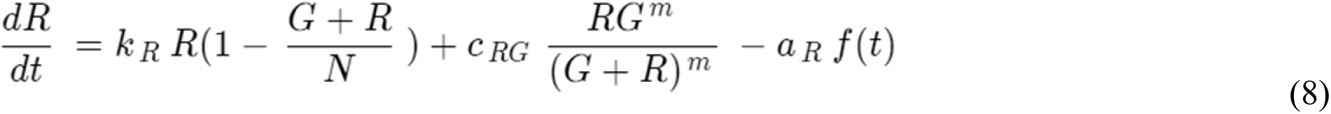

## Supplementary Figures

**Figure S1:**
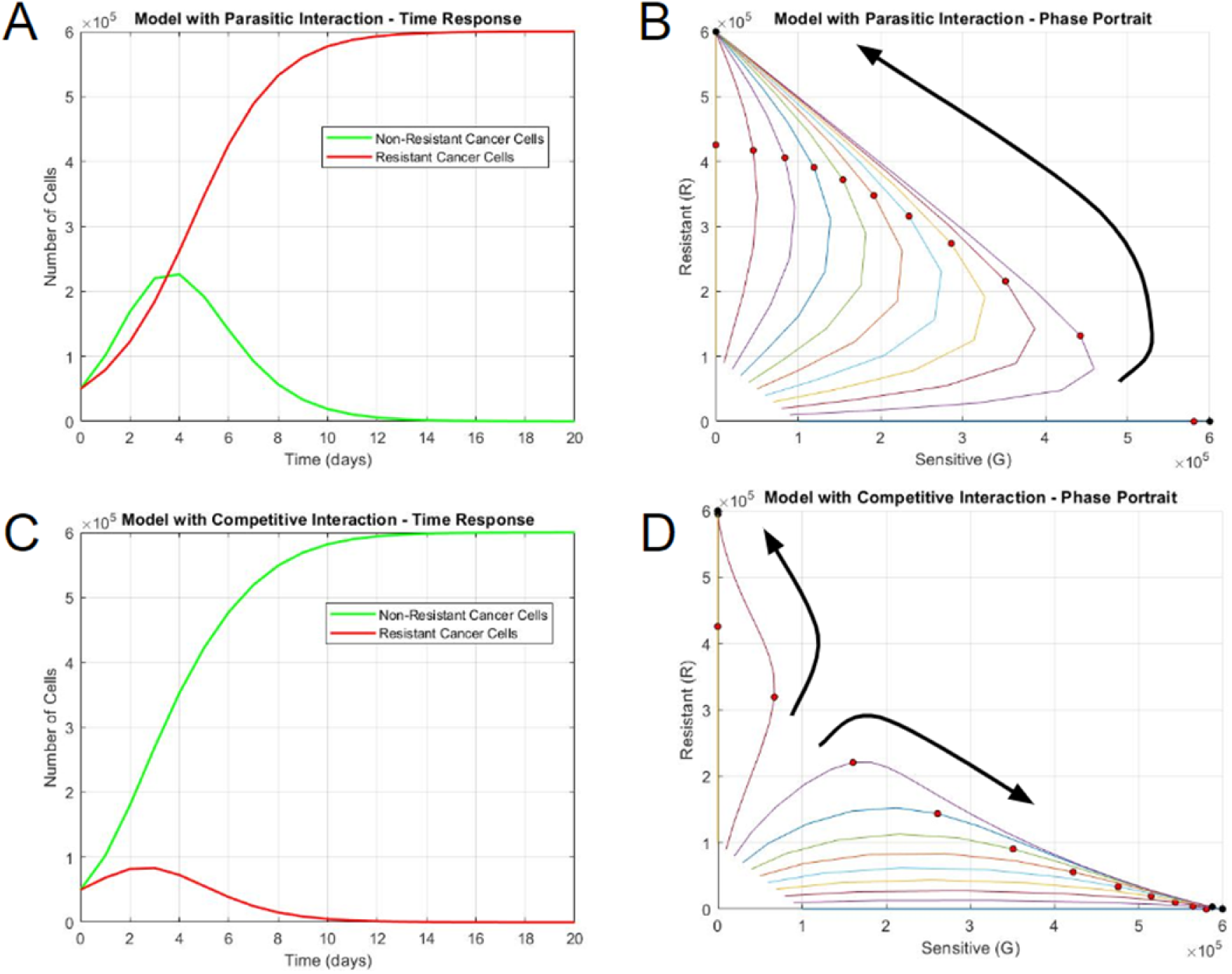
Long-term growth behaviors differentiate non-interacting and interacting cell populations. A) Time response and B) Phase portrait of logistic model with parasitic interaction. C) Time response and D) Phase portrait of logistic model with competitive interaction. In the phase portraits, the horizontal axis represents the number of sensitive DLBCL cells and the vertical axis represents the number of resistant DLBCL cells. Each curve in the phase portrait was created by altering the percentage of sensitive and resistant cells while keeping a constant initial population. The larger black arrows represent the general trajectories of population growth. The red circles represent the states on day 5, the black circle represent the steady state(s) at much longer time periods (day 100).

**Figure S2:**
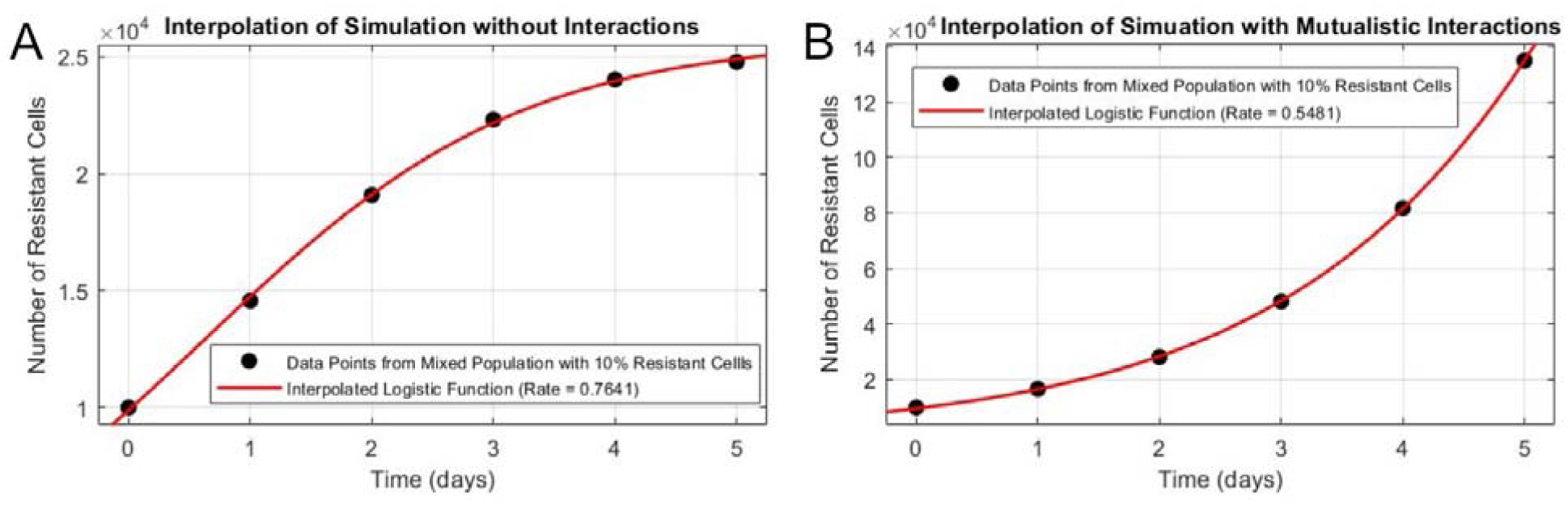
Example of rate constant interpolation from initial simulations. Sensitive and resistant cells from each simulation were separately fitted into a logistic equation to obtain rate constants (*k*). A) Non-interacting resistant cells in a 10% resistant cell culture demonstrate a rate constant of 0.7641 day^-1^, whereas B) Interacting resistant cells in a 10% resistant cell culture demonstrate a rate constant of 0.5481 day^-1^.

**Figure S3:**
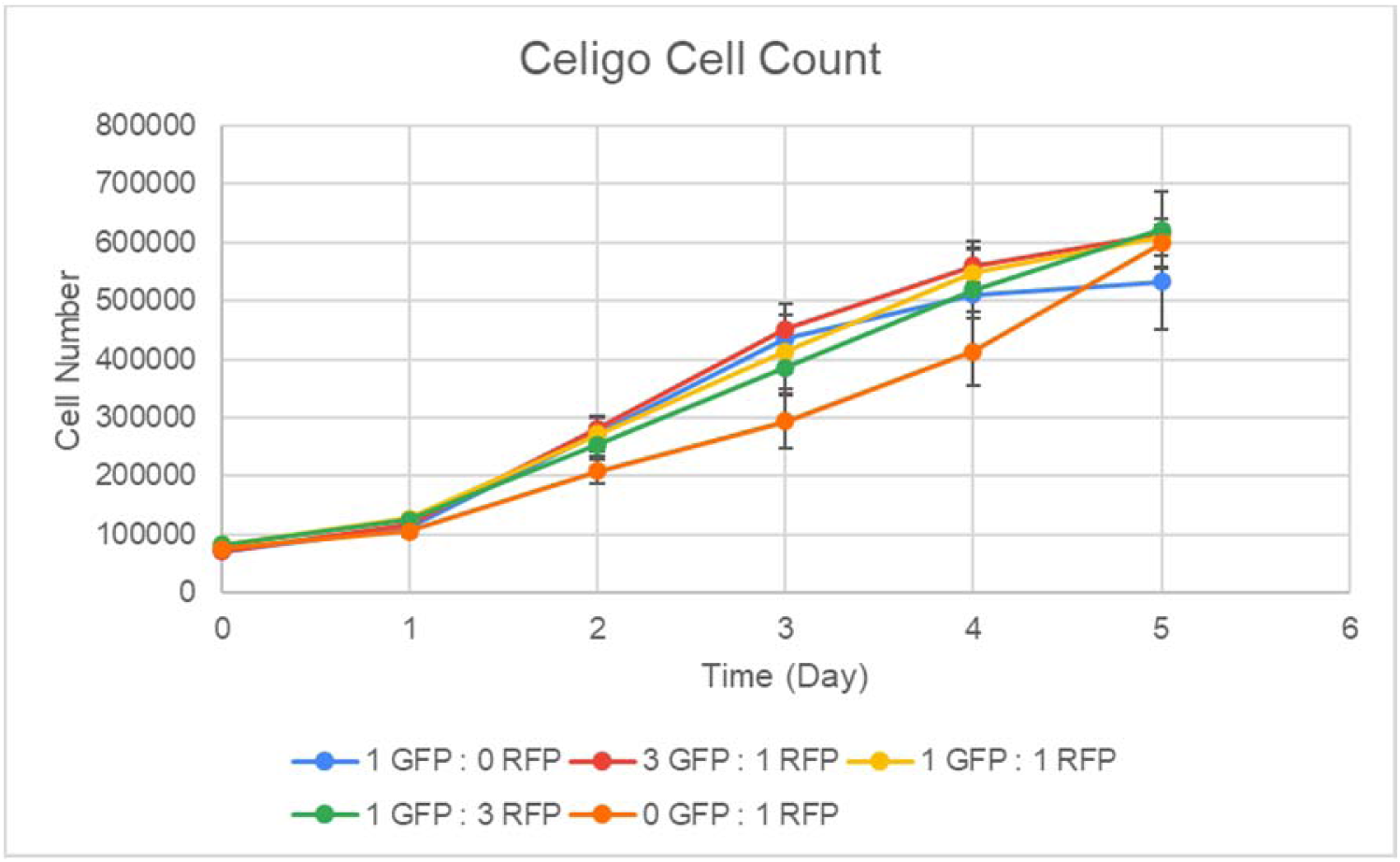
Total cell count of pure and mixed cultures from Celigo imaging cytometer. Cultures represent logistic growth.

**Figure S4:**
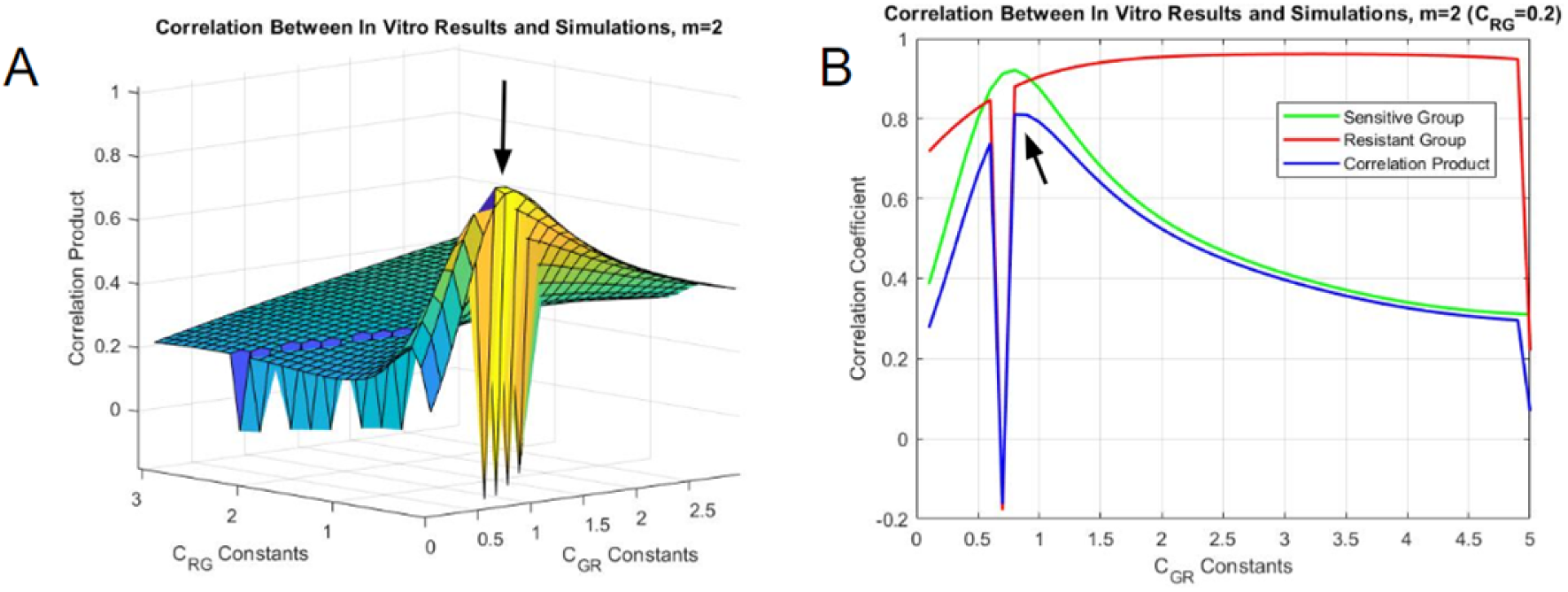
Optimization of the correlation product with **A)** 3-dimensional and **B)** 2-dimensional representations. The horizontal axes represent the range of tested interaction constants (C_GR_ and/or C_RG_). The vertical axis represents the correlation coefficients derived from correlating *in vitro* interpolated rate constants with rate constants from simulations using the indicated interaction coefficients. The correlation product is derived from multiplying correlation coefficients from the sensitive and resistant populations.

**Figure S5:**
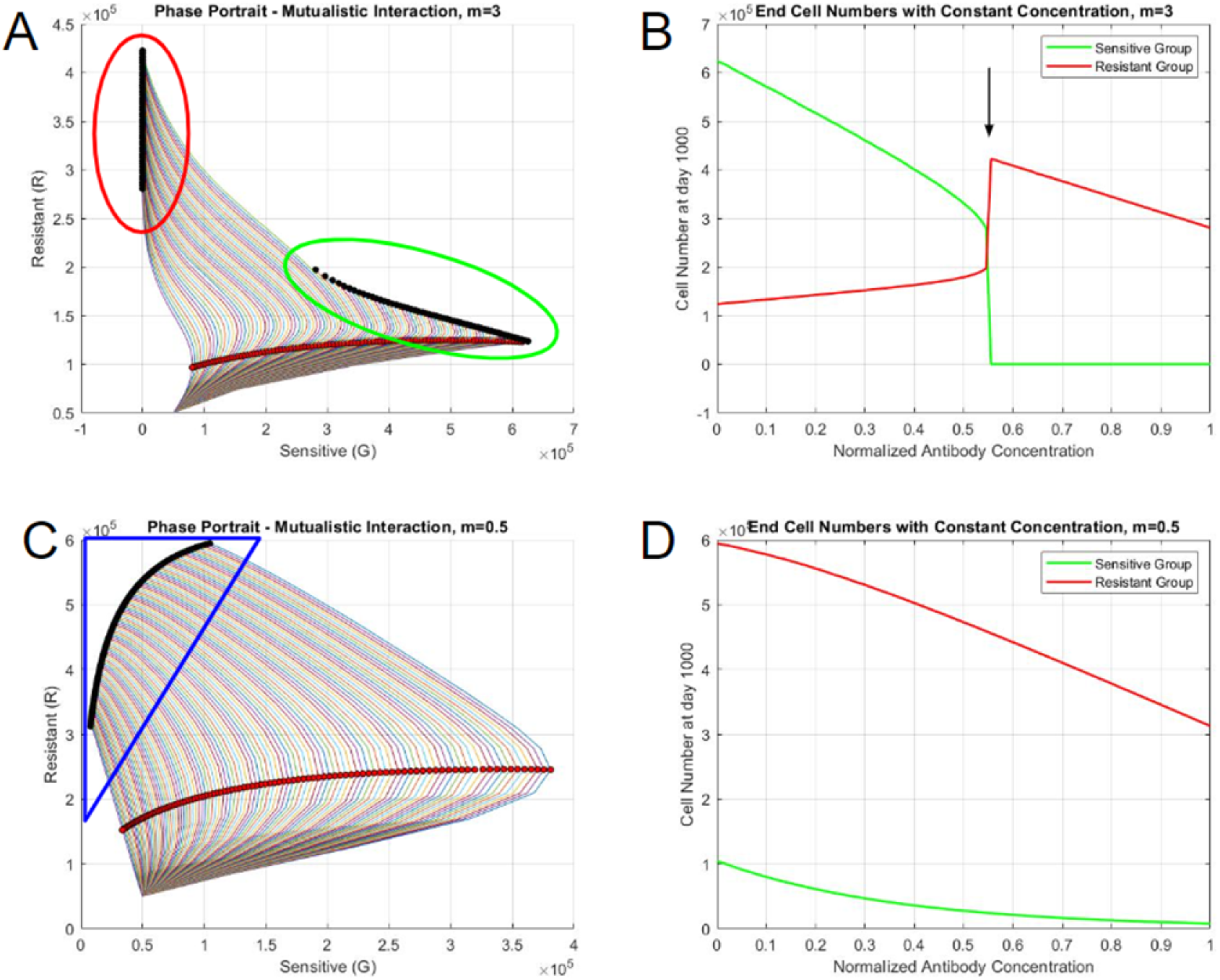
Steady state behaviors of systems with constant therapy. A) Phase portrait when m=3, B) End cell numbers when m=3, C) Phase portrait when m=0.5, D) End cell numbers when m=0.5. The red circles represent the states on day 5, the black circles represent the steady states. The outlines encircling sets of steady states represent distinct behaviors based on the concentration of therapy.

**Figure S6:**
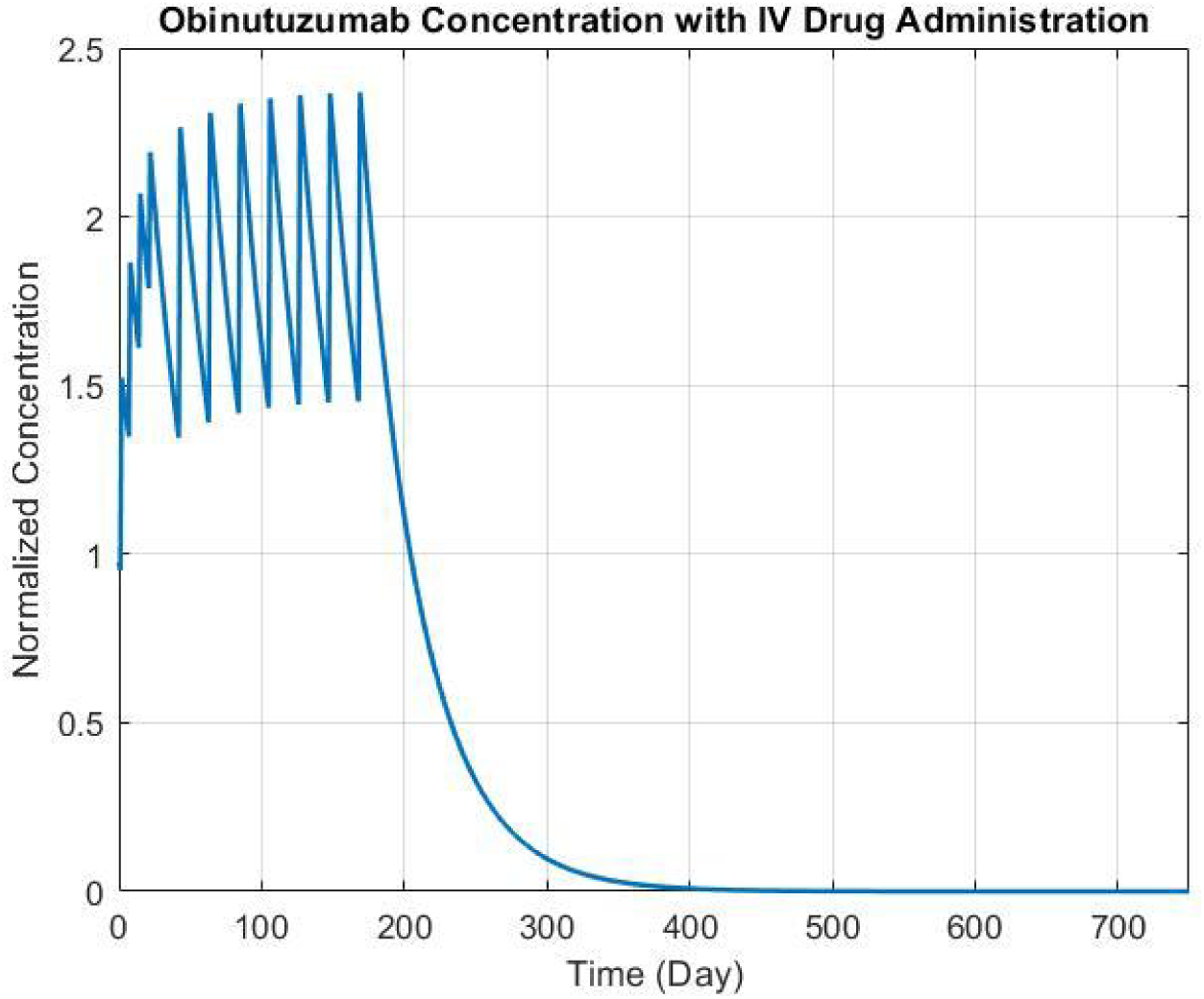
Obinutuzumab concentration over time based on therapy regimen. The vertical axis represents the concentration of therapy when the magnitude of the exponential function is set to one.

**Figure S7:**
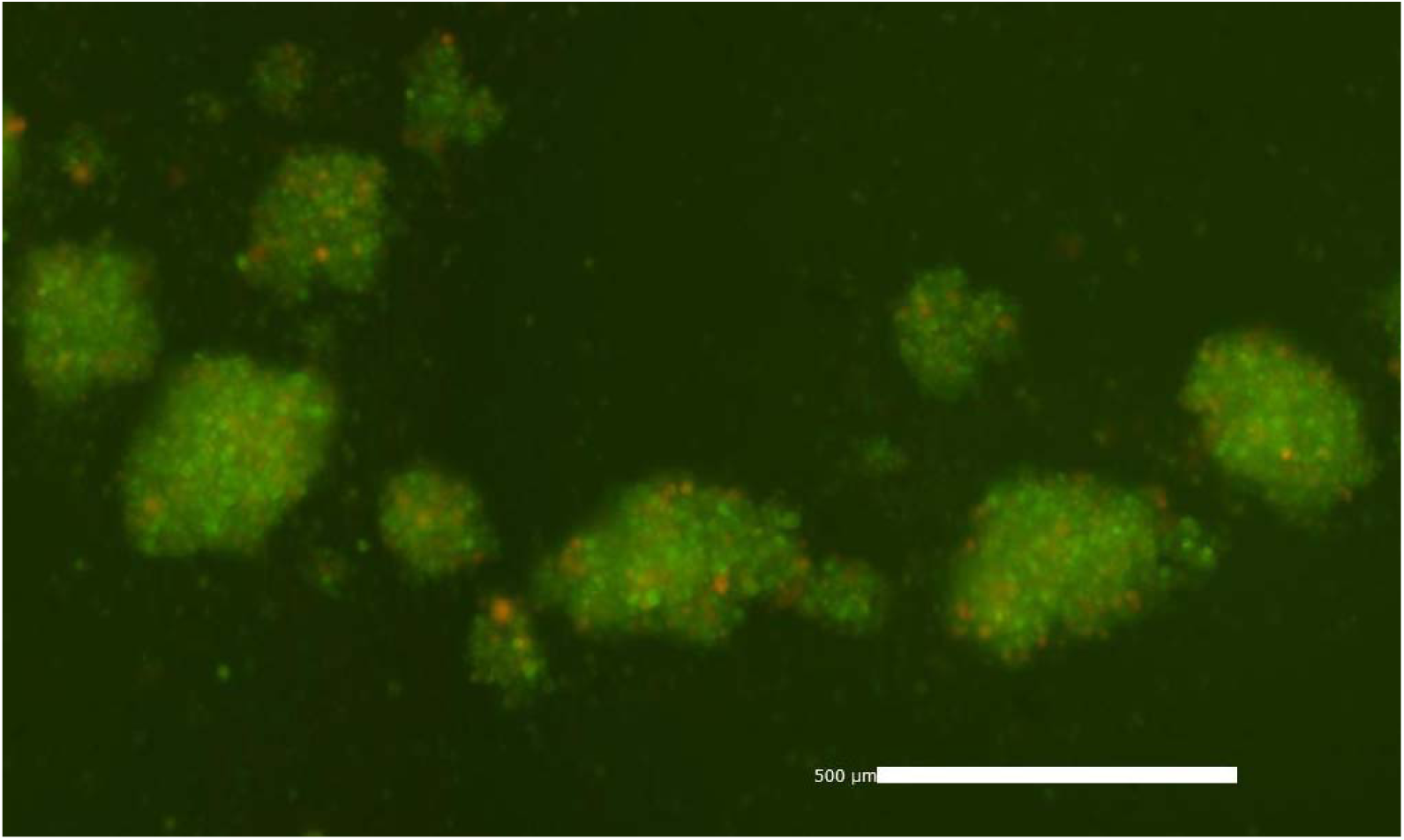
Aggregation of cells in mixed culture imaged with Celigo imaging cytometer - the aggregates contain SUDHL-4 (green) and SUDHL-4OR (red) cells.

## Supplementary Tables

**Table S1:** Interpolated constants from *in vitro* data.

Green Fluorescent Protein (GFP) Constants
|  | Initial Proportion of Cells |  |  |  |  |
| --- | --- | --- | --- | --- | --- |
|  | 1 GFP : 0 RFP | 3 GFP : 1 RFP | 1 GFP : 1 RFP | 1 GFP : 3 RFP | 0 GFP : 1 RFP |
| k (Rate Constant, day<sup>-1</sup>) | 1.069 | 1.133 | 1.191 | 1.065 | Not Applicable |
| CC (Carrying Capacity, number of cells) | 2.62E+05 | 2.72E+05 | 2.22E+05 | 1.81E+05 | Not Applicable |
| P(0) (Initial Population, Number of Cells) | 2.83E+04 | 2.38E+04 | 1.37E+04 | 8030 | Not Applicable |
| Adjusted R<sup>2</sup> | 0.9905 | 0.9934 | 0.9918 | 0.9985 | Not Applicable |

Red Fluorescent Protein (RFP) Constants
|  | Initial Proportion of Cells |  |  |  |  |
| --- | --- | --- | --- | --- | --- |
|  | 1 GFP : 0 RFP | 3 GFP : 1 RFP | 1 GFP : 1 RFP | 1 GFP : 3 RFP | 0 GFP : 1 RFP |
| k (Rate Constant, day<sup>-1</sup>) | Not Applicable | 0.5128 | 0.6045 | 0.596 | 0.4695 |
| CC (Carrying Capacity, number of cells) | Not Applicable | 6.63E+04 | 7.57E+04 | 1.33E+05 | 3.10E+05 |
| P(0) (Initial Population, Number of Cells) | Not Applicable | 5401 | 8126 | 14000 | 22300 |
| Adjusted R<sup>2</sup> | Not Applicable | 0.9886 | 0.973 | 0.9488 | 0.9663 |

**Table S2:** Correlation products and interaction constants based on exponent value.

| Exponent (m) | Correlation Product | Interaction Coefficients (day<sup>-1</sup> cell<sup>-m</sup>) |  |
| --- | --- | --- | --- |
|  |  | C<sub>GR</sub> | C<sub>RG</sub> |
| 0.5 | 0.8078 | 0.08 | 0.2 |
| 1 | 0.7943 | 0.27 | 0.15 |
| 2 | 0.8105 | 0.8 | 0.2 |
| 3 | 0.8447 | 2.3 | 0.2 |

**Table S3:** Constants based on clinical data.

938 Constants When the Interaction Exponent (m) is Equal to 2
|  | *In Vitro*<br>Constants | *In Vivo* Constants<br>(Slow Growth) | *In Vivo* Constants<br>(Fast Growth) |
| --- | --- | --- | --- |
| $k_G$ (Sensitive Cell Rate Constant, day<sup>-1</sup>) | 1.068 | 0.0239 | 0.0495 |
| $K_R$ (Resistant Cell Rate Constant, day<sup>-1</sup>) | 0.4695 | 0.0105 | 0.0218 |
| CC (Carrying Capacity, number of cells) | 6.00E+05 | 1.10E+12 | 1.10E+12 |
| $C_{GR}$ (Sensitive Cell Interaction Coefficient, day<sup>-1</sup>cell<sup>-2</sup>) | 0.8000 | 0.0179 | 0.0371 |
| $C_{RG}$ (Resistant Cell Interaction Coefficient, day<sup>-1</sup>cell<sup>-2</sup>) | 0.2000 | 0.0045 | 0.0093 |

Constants When the Interaction Exponent (m) is Equal to 1
|  | *In Vitro*<br>Constants | *In Vivo* Constants<br>(Slow Growth) | *In Vivo* Constants<br>(Fast Growth) |
| --- | --- | --- | --- |
| $k_G$ (Sensitive Cell Rate Constant, day<sup>-1</sup>) | 1.068 | 0.0239 | 0.0495 |
| $K_R$ (Resistant Cell Rate Constant, day<sup>-1</sup>) | 0.4695 | 0.0105 | 0.0218 |
| CC (Carrying Capacity, number of cells) | 6.00E+05 | 1.10E+12 | 1.10E+12 |
| $C_{GR}$ (Sensitive Cell Interaction Coefficient, day<sup>-1</sup>cell<sup>-1</sup>) | 0.2700 | 0.0060 | 0.0125 |
| $C_{RG}$ (Resistant Cell Interaction Coefficient, day<sup>-1</sup>cell<sup>-1</sup>) | 0.1500 | 0.0034 | 0.0070 |

